# Orchard management alters citrus root and rhizosphere microbiomes with functional consequences for plant performance

**DOI:** 10.64898/2026.04.14.717254

**Authors:** Nichole Ginnan, Riley Jones, Jessica Wu-Woods, Tariq Pervaiz, Ashraf El-kereamy, Vanessa E.T.M. Ashworth, M. Imran Hamid, Emma K. Dawson, Sarah L. Strauss, Jason Stajich, Philippe Rolshausen, M. Caroline Roper

## Abstract

Agricultural management practices act as ecological disturbances that can restructure soil and plant-associated microbial communities, but the functional consequences of these microbial shifts on crop performance remain poorly understood. Here, we examined how common orchard inputs, including wood mulch, glyphosate, and humic acid, affect citrus root and rhizosphere microbiomes and tree performance over a three-year field experiment. Mulch emerged as the dominant driver of microbiome structure, significantly altering bacterial and fungal community composition and increasing rhizosphere alpha diversity. Root microbiomes remained comparatively stable, suggesting stronger host selective forces within root tissues. Mulched rhizospheres were enriched with saprotrophic fungi and metabolically diverse bacteria, while non-mulched soils contained taxa typically associated with nutrient cycling, like *Rhizobium*, *Sphingomonas*, and *Nitrososphaera*. Interactions between mulch and glyphosate further reshaped bacterial communities and corresponded with reduced tree physiological performance, including photosynthesis rates. To verify whether these microbial shifts were contributing to these plant phenotype changes, we conducted a greenhouse experiment using field-derived soil microbiota. Active microbiota from mulch-treated soils reduced citrus seedling establishment and root growth relative to microbiota from non-mulched soils, whereas heat-killed controls eliminated these negative effects, demonstrating a causal relationship between management-induced microbiota changes and decreases in plant performance. In contrast, humic acid influenced plant growth primarily through direct abiotic effects rather than microbial community-level traits. Together, our results show that orchard management practices can restructure citrus microbiomes and generate community-level traits that influence plant performance, highlighting the importance of incorporating microbial ecology and microbiome information when designing and testing crop management strategies.

## Introduction

Growers are increasingly challenged to balance high productivity, rising input costs, and environmental impacts to maintain profitable and sustainable cropping systems [1]. Achieving this balance requires agricultural practices that work with, rather than against, the ecological processes that support plant growth. Historically, most management strategies were developed with limited to no consideration of soil microbial communities. However, it is now clear that soil-and plant-associated microbiomes play vital roles in regulating plant nutrition, mediating stress tolerance, and suppressing pathogens [2–7]. Agricultural inputs such as herbicides, organic amendments, and ground cover management alter plant and soil microbial communities [8]. Understanding how management practices influence microbiome composition and function is a critical step towards developing approaches to steer microbial communities to sustainably enhance plant performance.

Management practices can influence plant performance directly and indirectly by modifying microbiome functions and plant-microbe interactions. For example, field amendments and cover crops can alter plant root exudates, which impacts plant microbiome assembly [9–12]. From an ecological perspective, many agricultural inputs function as disturbances that alter microbial community composition, diversity, and functional potential [13, 14]. While the negative impacts of mechanical disturbances, like tillage, have been well characterized [15–17], the effects of commonly used herbicides and organic inputs on microbial community form and function are less well understood, particularly for perennial tree crops.

Glyphosate-based herbicides are widely used to manage weeds in agricultural systems. Glyphosate blocks the activity of 5-enolpyruvylskhikimate-3-phosphate synthase, an enzyme in the shikimate pathway, which is essential for the synthesis of important aromatic amino acids [18, 19]. In addition to its herbicidal activity, glyphosate can alter soil, plant, and animal gut microbial communities, as many bacteria and fungi also possess the targeted pathway [18, 20–22]. Previous studies have reported glyphosate-induced reductions in microbial diversity and plant growth promoting taxa, as well as increases in glyphosate-degrading microorganisms [20]. For example, glyphosate can decrease arbuscular mycorrhizal fungal colonization of roots [23], and increase abundance of putative pathogens, like *Fusarium* spp. [24, 25]. Because glyphosate is commonly applied alongside fertilizers and other amendments, interactions among these inputs may further complicate their effects on soil microbial communities and plant performance.

Organic amendments also influence soil properties and microbial communities [26, 27]. Wood mulch is often applied in perennial cropping systems to suppress weeds and retain soil moisture [28, 29]. Mulch can also alter soil nutrient availability and has been associated with increased microbial diversity and shifts in rhizosphere community composition [30]. Similarly, humic acid, an organic component of soil organic matter, can stimulate plant growth and root development, through direct modulation of plant physiology [31, 32] and indirect microbial community alterations, such as enrichment of pathogen-suppressive bacteria [33]. Furthermore, organic amendments, such as earthworm cast, can decrease glyphosate phytotoxicity when applied in combination [34], again highlighting the need to study the combined effects of herbicide and organic amendments on plant-associated microbiota.

Citrus is an important global tree crop and has emerged as a model system for studying plant-microbiota interactions in perennial agriculture [35]. Citrus trees harbor diverse microbial communities across root, rhizosphere, and aboveground tissues [36]. These microbiomes vary across plant compartments, host genotypes, and geographic locations, while still maintaining a core set of microbial taxa [36–40]. Shifts in citrus-associated microbiomes have been linked to plant developmental stage, disease progression, and seasonality [41–43]. Moreover, the citrus microbiome is a reservoir of microbial taxa with potential to help control citrus pathogens, understand disease holistically, and promote growth and fruit quality [44–49].

Despite the growing recognition that management practices influence crop microbiomes, it remains difficult to determine whether these changes translate into meaningful functional effects on plant performance. Microbial kingdoms (bacteria, fungi, archaea) often have different responses, and while studies have revealed associations between management practices, microbiome composition, and plant phenotypes, experimental validation is needed to establish causal relationships [50].

In this study, we investigated how common orchard management practices influenced citrus microbiomes and tree performance over three years. Field plots received combinations of wood mulch, glyphosate, and humic acid applications in a factorial design. We first evaluated the effects of these treatments on tree performance, weed management, and root and rhizosphere bacterial and fungal communities in the field. Then, to test whether management-driven microbiome shifts persist and translate into functional consequences for plant growth, we conducted a greenhouse experiment where soil microbiota from the field plots were inoculated into sterile substrate and used to propagate citrus rootstock seedlings. By comparing active microbial inocula with heat-killed controls, we distinguished microbial effects from abiotic soil effects. Together, these experiments determined that common orchard management practices do indeed alter microbiome community-level traits in ways that influence citrus phenotypes.

## Materials and Methods

### Field experiment site and design

The approximately three-year (April 2021– August 2024) field experiment was conducted at the Lindcove Research and Extension Center (LREC) in Exeter, California, USA. The study included eight treatment plots (16 trees/plot) with ‘Tango’ mandarin orange (*Citrus reticulata* Blanco) trees grafted on Carrizo rootstock [*Citrus sinensis* (L.) Osbeck x *Poncirus trifoliata* (L.) Rafinesque] planted in 2011. Plots were established in a full factorial design crossing three management factors: wood mulch, glyphosate (herbicide), and humic acid (HA) (each applied or not applied), resulting in eight treatment combinations (Table 1). The combination in which none of the three factors were applied served as the untreated control.

**Table 1.** Field application treatment groups.

| Treatment Group ID | Mulch | Glyphosate | Humic Acid |
| --- | --- | --- | --- |
| CCC | – | – | – |
| CCH | – | – | yes |
| CGC | – | yes | – |
| CGH | – | yes | yes |
| MCC | yes | – | – |
| MCH | yes | – | yes |
| MGC | yes | yes | – |
| MGH | yes | yes | yes |

The wood mulch initially consisted of chipped almond tree material sourced from a local nursery (Visalia, CA, USA). A full mulch application was completed in April 2021 at 10 tons/acre. Mulch was applied across the whole plot, including between rows and beneath canopies to a depth of 7-8 cm (Supplemental Fig. S1). Mulch from chipped citrus trees originating from the LREC was reapplied in April 2023 at approximately 8 tons/acre.

Glyphosate (Roundup original, EPA# 524-445, Bayer Crop Sciences, NC, USA) was applied in March, May, and August, at 16 oz per 50 gallons of water. HA (Growth Boost, SoilBiotics, Kankakee, IL, USA) was applied under the tree canopy around the trunk at 5 gallons/acre monthly from April to October. After application sprinklers were turned on for 30 minutes to facilitate product absorption into the soil and roots.

All treatment plots received identical standard management practices. Irrigation was applied at two-week intervals for 48 hours per watering event, totaling approximately 3 acre-feet of water per year. Twenty units of calcium ammonium nitrate (17-0-0; CAN 17) was applied monthly from April through August.

### Data collection and field sampling

Photosynthetic measurements were collected each year in June (2022, 2023, 2024) on fully expanded sun-exposed leaves between 11:00 AM and 12:30 PM, using a LI-6800 portable photosynthesis system (Li-Cor, Inc., Lincoln, NE, USA). The values of net photosynthetic activities including stomatal conductance (gsw), carbon assimilation (photosynthetic) rate (A), transpiration rate (E), and intercellular CO_2_ concentration (Ci), were recorded. Measurements were repeated on four leaves (technical replicates) on the west side of the trees. Six replicate trees were sampled per treatment plot each year for a total of n=192.

Fruit was harvested between February 28th and March 5th of 2022 and 2023. The fruits from each tree (8 trees per treatment group, n=64) were transported in bins to an on-site pack line where total fruit weight and fruit counts were collected. Average fruit weight was calculated by dividing the total weight by the number of fruits.

Weed aboveground biomass was collected in August 2023 and 2024 by dividing each plot by tree rows and collecting weeds from three replicate rows per treatment plot using a sickle. Weed fresh weight was measured. Then, samples were dried for two months under ambient conditions and dry weight was measured.

### Microbiome sample collection and processing

Each year (05/24/2022, 05/30/2023, 06/04/2024) root and rhizosphere samples were collected (n= 96 tree samples; 4 biological replicates x 8 treatment groups x 3 years) for microbiome analyses. Root samples were collected near the irrigation line of each tree by removing the top 5 cm layer and digging into the root zone 15-20 cm and collecting ∼10 g of roots in 50 ml falcon tubes. Roots were washed twice with 25 ml of sterile diH_2_O by vigorously shaking the falcon tubes. Roots and liquids were then separated, flash frozen in liquid nitrogen, and stored at - 80°C.

Root tissue was lyophilized in a FreeZone 2.5 L benchtop freeze dryer (Labconco, Kansas City, USA) for 72 h. Lyophilized tissue was ground into a powder on a MM 400 mill (Retsch, Haan, Germany) in 35 mL stainless steel grinding jars, each containing a 20 mm stainless steel ball, at 25 oscillations/sec delivered in 30 second increments.

### Microbial DNA extractions, library preparations and amplicon sequencing

Microbial DNA was extracted and purified using a high-throughput workflow implemented on a QIAcube HT robotic instrument (QIAGEN, Hilden, Germany), for more details see Supplemental Methods S1. Amplicon sequencing libraries were prepared following Earth Microbiome Project protocol [51, 52], see Supplemental Methods S2 for a full description.

Library sequencing was completed by the UC Riverside Genomics Core Facility on an Illumina NextSeq (2000 P1 600 cycle kit). NextSeq 2000 runs provide up to 100 million reads per run and ∼450 samples were multiplexed per run for an average of 222,222 reads per sample.

### Amplicon sequence processing

Raw Illumina sequences were demultiplexed, with zero index barcode mismatches allowed, and converted to FASTQ format using BCL Convert v3.8.4. Adaptors and primer sequences were removed using trimmomatic [53] with standard settings and cutadapt [54] with a minimum 5-base overlap, maximum 15% error rate when matching primers, and used -–nextseq-trim=20 to trim polyX tails and low-quality ends. FastQC [55] was used to verify read quality and check for adapter sequences. Additional quality control and reads processing was completed using the DADA2 pipeline [56], see full workflow in Supplemental Methods S3.

Post-processing reads associated with plant contamination were removed from the ASV and taxa tables. Then, a threshold that discarded any ASVs that did not appear in at least five samples and had at least 20 reads was applied. ASVs with less than 100 total reads across all samples were removed to reduce rare ASVs and potential contaminants. Then, samples with <1000 reads were removed. The ZymoBIOMICS Microbial Community Standard (positive control) samples were evaluated and indicated that all taxa were present and contaminant ASVs represented less than 1% of the sample community. The positive and negative control samples were removed for downstream analyses.

### Microbial community analyses

The cleaned and prepared microbiome data, as phyloseq objects, were loaded into R (v.4.5.0) [57, 58]. Bacterial and fungal Shannon and inverse Simpson (InvSimpson) indices were calculated using phyloseq::estimate_richness() and were fit to mixed-effects models using the lme4 (v.1.1-37) package [59], when possible, or a fixed-effects model if overfitting of the mixed-effects model occurred. Data was visually assessed for outliers, which were removed. Removing the outliers did not impact the interpretation of any of the results, only improved the model fit. Initial formulas used for the mixed-effects or linear models, respectively, include:

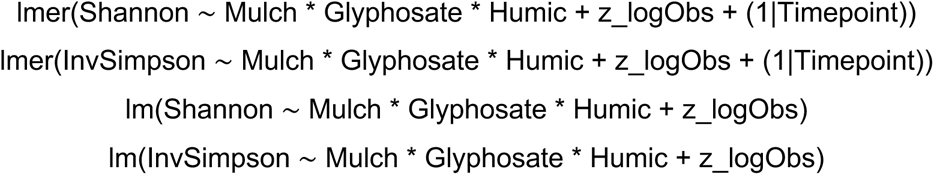

where the random effect “Timepoint” represents the sample year, and “z_logObs” represents the z-scores of the log of the total observed sequence in that sample, which helps control for variation in sample sequencing depth. Model reduction proceeded by sequentially removing non-significant higher-order interactions to obtain the most simplified model. A type III ANOVA was performed using stats::anova(), to test for significance, followed by pairwise comparisons and significance tests using the emmeans (v.1.10.2) package [60] with FDR adjustments. All final models and ANOVA results are provided in Supplemental Table S1. Data visualizations were generated using ggplot2 (v.3.5.1) [61].

Bacterial community beta diversity was assessed with a constrained analysis of principal coordinates of centred-log-ratio-transformed counts, using phyloseq::ordinate, method = ‘CAP’, and distance = ‘Euclidean’ with the following formula:

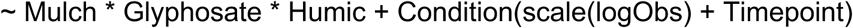

Model reduction proceeded by sequentially removing non-significant higher-order interactions to obtain the most simplified model. An ANOVA-like permutation test was used to test for model significance using the vegan::anova.cca() function [62]. The anova.cca by = ‘term’ option was used to determine the significance of the Mulch, Glyphosate, and Humic variables separately. CAP1 and CAP2 axes were plotted using ggplot2, including 95% confidence interval ellipse (stat_ellipse()).

A bacterial and fungal genus-level differential abundance analysis was completed using ALDEx2 v.1.41.0 [63, 64]. Briefly, this program uses centered log-ratio (CLR) transformation of raw counts and Monte-Carlo sampling (mc.samples=200) to estimate the probability that a taxon is enriched. Then, we used ALDEx2::aldex.ttest(), which employs a Wilcoxon test and Benjamini-Hochberg correction to test for significance. Results for all tested genres are listed in Supplemental Table S2. Effect size (magnitude of difference between two groups) was calculated using ALDEx2::aldex.effect(). Volcano plots were created using ggplot2 [61].

### Field measurements analyses

Raw field data is available in Supplemental Table S3. All field measurement data were analyzed in R (v.4.4.1) [57, 58]. Field measurements were fit to mixed-effects models as described above. Initial formula used for the mixed effects:

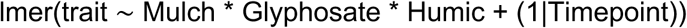

where the random effect “Timepoint” represents the sample year. Model reduction, statistical tests, and visualizations were completed using methods outlined above. All final models and ANOVA results are provided in Supplemental Table S1.

### Soil microbiota functional greenhouse experiment

A greenhouse experiment was conducted to examine plant responses to field-derived soil microbiota. Field soil was collected from each of the eight field treatment plots at LREC (Exeter, California, USA) on October 8, 2025. At this time, the mulch, glyphosate, and HA treatments had not been actively applied in approximately 14 months (field applications concluded in August 2024), but the standard fertilizer and irrigation continued. To collect the soil, three trees within each plot were randomly selected for soil sampling, and soil collected from these trees was pooled to generate one representative sample per plot. For each tree, soil was collected from a single side approximately 0.5 meter from the trunk. The top 5 cm of soil was removed, and soil was collected to a depth of 25 cm. Approximately 0.5 L of soil was obtained per tree (≈1.5 L total per plot). Clean gloves were worn, and shovels were sterilized with bleach between plots. Soil was transported back to the lab at room temperature and then stored at 4 °C.

Each of the eight soils collected had a portion separated and autoclaved twice for a 45-minute liquid cycle (cooling completely between cycles) to create “heat-killed” microbiota controls. This created a total of 16 treatment groups (8 active, 8 heat-killed).

Clean Carrizo rootstock seeds were purchased from Lyn Citrus Seed, Inc (Arvin, CA, USA) in September 2025. Prior to planting, seeds were soaked in sterile water at 28°C at 180 rpms for 24 hours, to split seed coats. Seed coats were peeled to increase germination uniformity and planted the same day.

The greenhouse experiment was initiated on November 7, 2025. Double autoclaved 70:30 (v/v) coco coir (#766310389395, EnvironSoil, Irvine, CA) to perlite (#100552701, Vigoro, Lake Forest, IL) soil-like substrate was prepared. Each of the 16 soil microbiota treatments were homogenized in individual batches with the sterile soil-like substrate with 10% field-collected soil and 90% sterile substrate (v/v) [65]. This method has been shown to effectively transfer microbiota, while reducing effects of soil structure and nutrients. Sterile plastic cone pots (#CN-SS-SC-07B, Stuewe & Sons, Inc., Tangent, OR) were filled with 150 ml of field soil inoculated substrate. One seed was planted per pot, approximately 20 mm deep, covered, and watered. Treatment pots (n=256, 16 biological replicates per treatment group) were set up in a randomized block design. Pots were maintained under ambient greenhouse conditions. Plants were uniformly watered two times per week.

After 2.5 weeks, pots with multiple seedlings emerged were thinned by gently removing the additional seedling(s). In the case of multiple seedlings, the largest was left because they are more likely to be nuclear clones [66], whereas smaller seedlings from the same seed are likely to be zygotic. Pots that did not have any seedlings emerge were noted and a seedling that was germinated in extra sterile substrate was transplanted into the pot.

After 14 weeks, the experiment concluded with measuring shoot height from the top of the soil to the top of the stem and recording seedling establishment. Then, plants were gently pulled from the soil and soil adhering to the roots was shaken or pulled off. Whole plants were placed in paper envelopes and air-dried at room temperature for six days and then dried in an oven at 60°C for 24 hours. After drying, dried roots and shoot mass were measured. Raw greenhouse experiment data is available in Supplemental Table S4.

### Greenhouse experiment analysis

All greenhouse data were analyzed in R (v4.5.0) [57, 58]. Plant establishment was analyzed as a binary response using a generalized linear model with a binomial error distribution and logit link. Firth’s bias-reduced logistic regression was implemented using the brglm2 package v1.0.1 [67, 68]. Fixed effects included mulch, glyphosate, and HA and their interactions. Significance of model terms was evaluated using a Likelihood Ratio Test. Estimated marginal means were calculated using the emmeans [60] package, and pairwise comparisons among treatment combinations were conducted with false discovery rate (FDR) adjustment for multiple testing.

Plant growth traits (dry root biomass, dry shoot biomass, and shoot height) were fit to mixed-effects models as described in the microbial community analysis section. Initial formula used for the mixed effects:

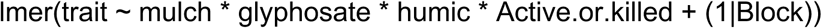

where the random effect “Block” controls for variation in greenhouse conditions across experimental blocks and “Active.or.killed” represents whether the microbiota are active or autoclaved (heat-killed). Model reduction, statistical tests, and visualizations were completed using methods outlined above. All final models and results are provided in Supplemental Table S1.

## Results

### Field applications primarily impacted rare bacterial, but dominant fungal rhizosphere taxa

To assess bacterial and fungal alpha diversity responses to mulch, glyphosate, and humic acid (HA), we calculated the Shannon and inverse Simpson (invSimpson) Indices. Both metrics account for richness and evenness, but differ in sensitivity to rare versus dominant taxa, respectively. Treatment effects were evaluated using mixed-effects models controlling for year and sequencing depth (Supplemental Table S1).

Mulch significantly increased rhizosphere bacterial Shannon index (Fig. 1a, ANOVA, F_1,77_=8.49, P=0.005) and had significant pairwise interactions across treatment groups with bacterial rhizospheres that received mulch, glyphosate, and HA in combination having the greatest Shannon index (Fig 1a). Fungal rhizosphere Shannon and bacterial rhizosphere invSimpson indices were not significantly impacted by treatments (Fig 1b-c). However, mulch significantly increased fungal rhizosphere invSimpson Index (Fig. 1d, ANOVA, F_1,78_=5.87, P=0.018), indicating that applications primarily influenced rare bacterial, but dominant fungal rhizosphere taxa.

**Figure 1.**
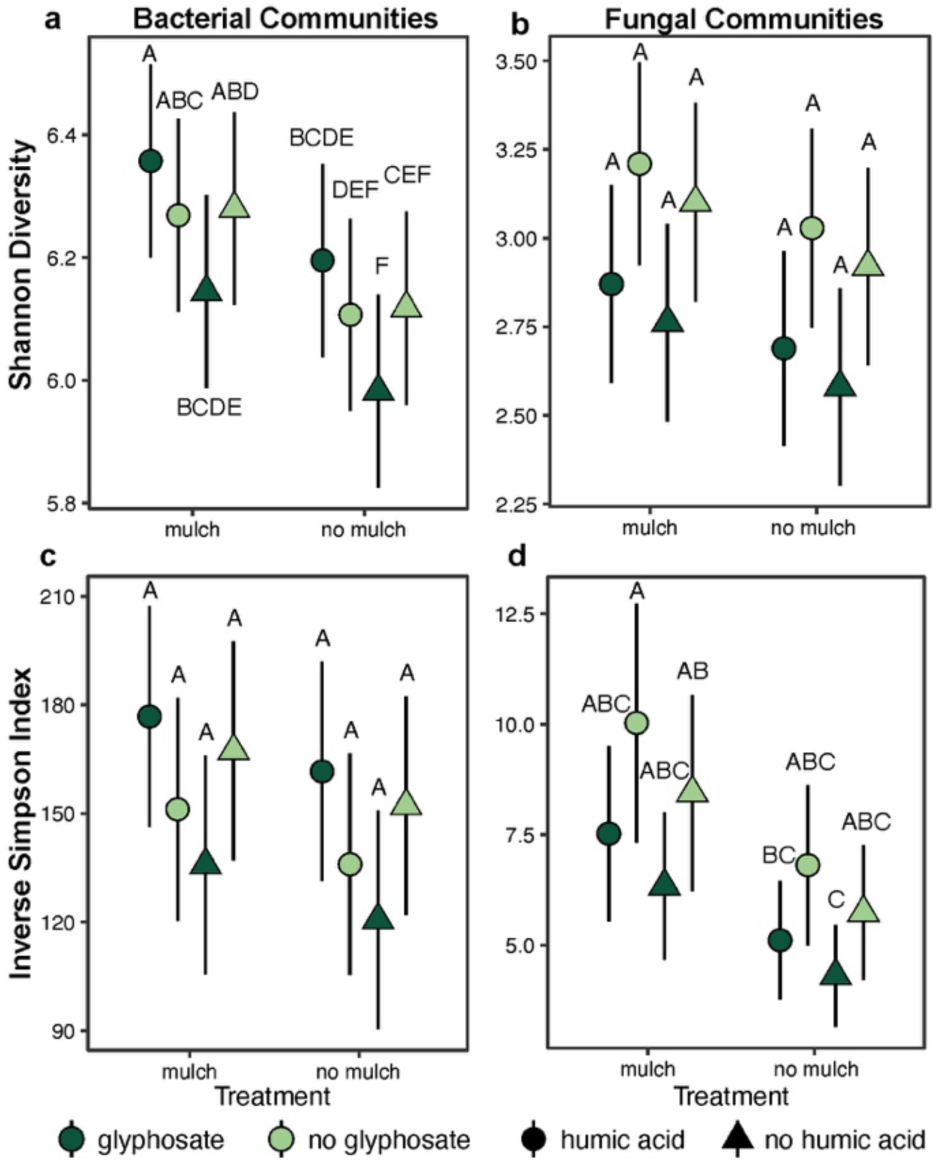
Mulch significantly increased rhizosphere bacterial and fungal alpha diversity. For the bacterial Shannon Index (a), fungal Shannon index (b), bacterial inverse Simpson index (c), and fungal inverse Simpson index (d) plots, points are estimated marginal means of biological replicates, error bars represent standard error, and letters indicate pairwise group differences based on ANOVA followed by a two-tailed Tukey post hoc test and FDR correction (P < 0.05). These analyses include n = 95 bacterial and n = 95 fungal rhizosphere microbiome samples. Light green represents samples from plots that were not treated with glyphosate and dark green represents samples from glyphosate treated plots. Circles indicate humic acid treated samples and triangles are samples that were not treated with humic acid. For example, a light green triangle indicates no glyphosate and no humic acid.

In roots, bacterial Shannon index was significantly impacted by mulch and glyphosate interaction (ANOVA, F_1,77_=4.11, P=0.046), but pairwise interactions were not significant, likely due to high data variability (Supplemental Fig. 1a, Supplemental Table S1). Bacterial and fungal invSimpson and fungal Shannon indices were not significantly impacted by the field treatments (Supplementary Fig. 1b-d), suggesting that rhizosphere communities are more responsive to management than root microbiomes.

### Mulch application was the major driver of root and rhizosphere microbiome composition

To test the effects of mulch, glyphosate, and HA on microbial community composition, we performed constrained ordinations on Euclidean distances, controlling for variation across years and sequencing depth (Supplemental Table S1). Field applications significantly affect bacterial and fungal community composition across roots and rhizosphere (Fig 2; Table 2). The bacterial rhizosphere community was the most sensitive, with treatments explaining 8.2% of the community variation (Fig 2; Table 2).

**Figure 2.**
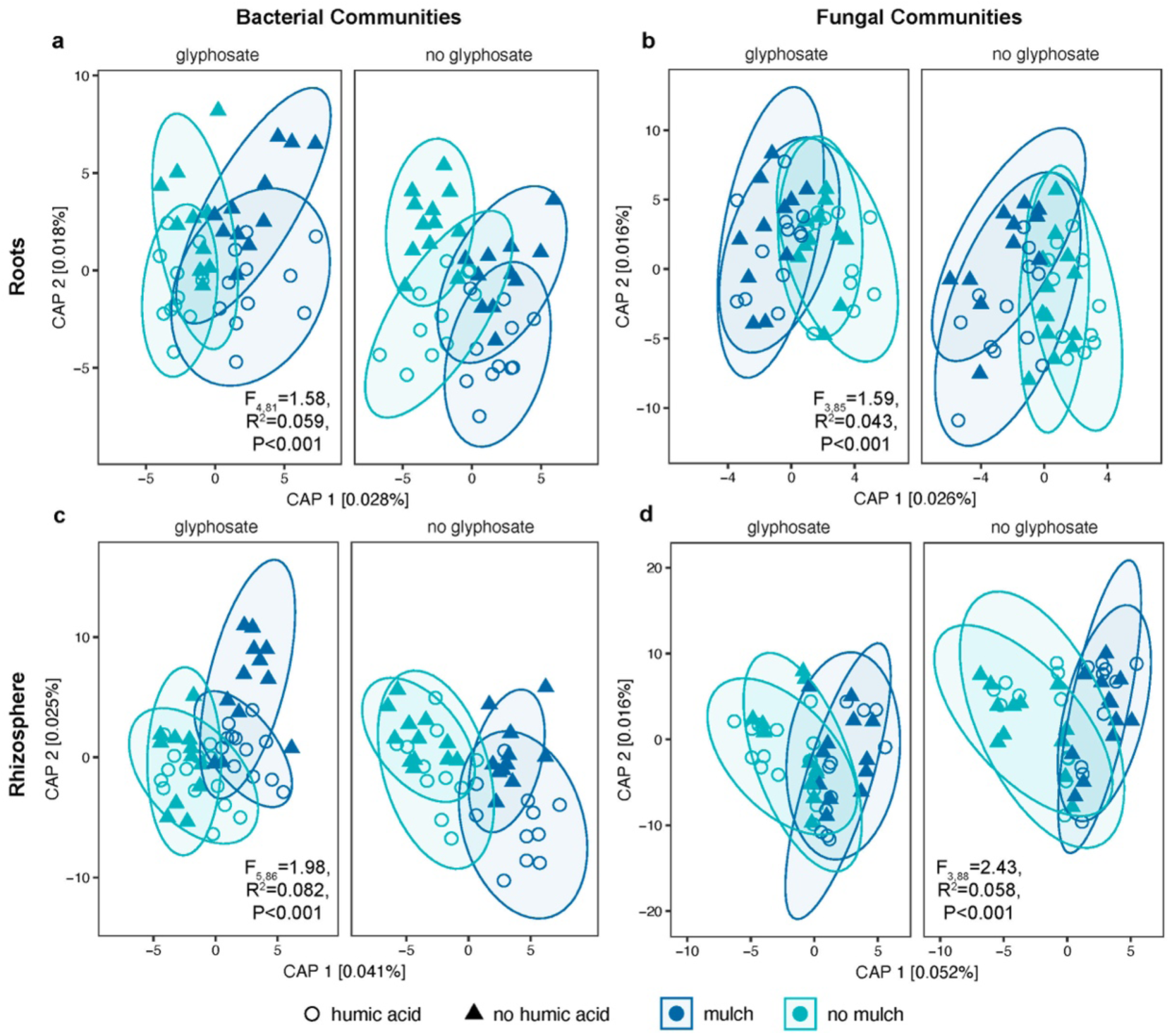
Field applications significantly impacted bacterial and fungal community composition. For the bacterial root (a), fungal root (b), bacterial rhizosphere (c), and fungal rhizosphere (d) plots are constrained ordinations by mulch, glyphosate, and humic acid with the full model ANOVA-like permutation test results indicated on each plot. Ellipses indicate 95% confidence intervals. These analyses include n = 95 bacterial and fungal rhizosphere and n=89 bacterial and n=92 fungal root microbiome samples.

**Table 2.** Canonical Analysis of Principal coordinates (CAP) ANOVA-like permutation test results, including the statistics for each term and interactions retained in the reduced model. Symbols correspond to significance level based on P-value.

| *Compartment* | *Community* | *Model Term* | *Df* | *Variance* | *F* | *P* | *Symbol* |
| --- | --- | --- | --- | --- | --- | --- | --- |
| Root | Bacteria | Full model | 4 | 208.13 | 1.5803 | ≤0.001 | *** |
|  |  | Mulch | 1 | 80.03 | 2.4306 | ≤0.001 | *** |
|  |  | Glyphosate | 1 | 39.99 | 1.2146 | 0.085 | . |
|  |  | Humic | 1 | 49.04 | 1.4892 | 0.012 | * |
|  |  | Mulch:Glyphosate | 1 | 39.07 | 1.1866 | 0.089 | . |
|  |  | *Residual* | 81 | 2667.1 |  |  |  |
| Rhizosphere | Bacteria | Full model | 5 | 585.1 | 1.9817 | ≤0.001 | *** |
|  |  | Mulch | 1 | 226.8 | 3.8411 | ≤0.001 | *** |
|  |  | Glyphosate | 1 | 70.2 | 1.1892 | 0.099 | . |
|  |  | Humic | 1 | 101.4 | 1.717 | 0.002 | ** |
|  |  | Mulch:Glyphosate | 1 | 99.1 | 1.6775 | ≤0.001 | *** |
|  |  | Mulch:Humic | 1 | 87.6 | 1.4839 | 0.014 | * |
|  |  | *Residual* | 86 | 5078.6 |  |  |  |
| Root | Fungi | Full model | 3 | 55.37 | 1.5893 | ≤0.001 | *** |
|  |  | Mulch | 1 | 25.41 | 2.1879 | 0.004 | ** |
|  |  | Glyphosate | 1 | 17.6 | 1.5152 | 0.042 | * |
|  |  | Humic | 1 | 12.37 | 1.0649 | 0.277 |  |
|  |  | *Residual* | 85 | 987.15 |  |  |  |
| Rhizosphere | Fungi | Full model | 3 | 176.26 | 2.4285 | ≤0.001 | *** |
|  |  | Mulch | 1 | 119.21 | 4.9277 | ≤0.001 | *** |
|  |  | Glyphosate | 1 | 35.87 | 1.4826 | 0.061 | . |
|  |  | Humic | 1 | 21.18 | 0.8753 | 0.591 |  |
|  |  | *Residual* | 88 | 2128.95 |  |  |  |

Across all four microbial communities examined, mulch application was the strongest predictor of microbial community composition (Fig. 2, Table 2). Glyphosate exerted detectable effects on community structure, ranging from marginally significant impacts in rhizosphere bacteria, rhizosphere fungi, and root bacteria to significant effects in root fungal communities (Table 2). HA only affected the bacterial, not fungal communities. Several interaction terms contributed to bacterial community variation, including mulch x glyphosate interaction, which significantly shaped rhizosphere bacterial composition and showed a marginal effect in root bacterial, and mulch x HA interaction, which significantly affected rhizosphere bacterial communities. These results indicate that field applications significantly reshape root and rhizosphere microbiome structure.

### Field applications significantly impacted tree productivity and weed biomass

Management-driven shifts in tree productivity and weed biomass were evaluated using mixed-effects models that included all sampling years, with year treated as a random effect to account for interannual variation (Supplemental Table S1). Across all models, HA was not a significant predictor of trait variation and was therefore excluded from the final analyses.

Fruit weight was significantly reduced by glyphosate application when applied alone, but this negative effect was reversed when glyphosate was combined with mulch, resulting in a significant increase in fruit weight relative to glyphosate-only plots (Fig. 3a; ANOVA, mulch x glyphosate, F_1,123_=4.204, P=0.042). Carbon assimilation rate declined significantly in response to mulch and was further reduced by glyphosate treatment, having an additive effect (Fig. 3b; ANOVA, mulch: F_1,186_=21.599, P=6.35×10^-06^; glyphosate: F_1,186_=7.049, P=0.009). Similarly, transpiration was significantly decreased under mulch and glyphosate treatment and these variables had an additive effect (Fig. 3c; ANOVA, mulch: F_1,186_=10.059, P=0.0018; glyphosate: F_1,186_=12.595, P=0.0005). Although not quantitatively assessed, chlorosis (leaf yellowing) was also observed in the canopy of mulch-treated trees (Supplemental Fig. S3).

**Figure 3.**
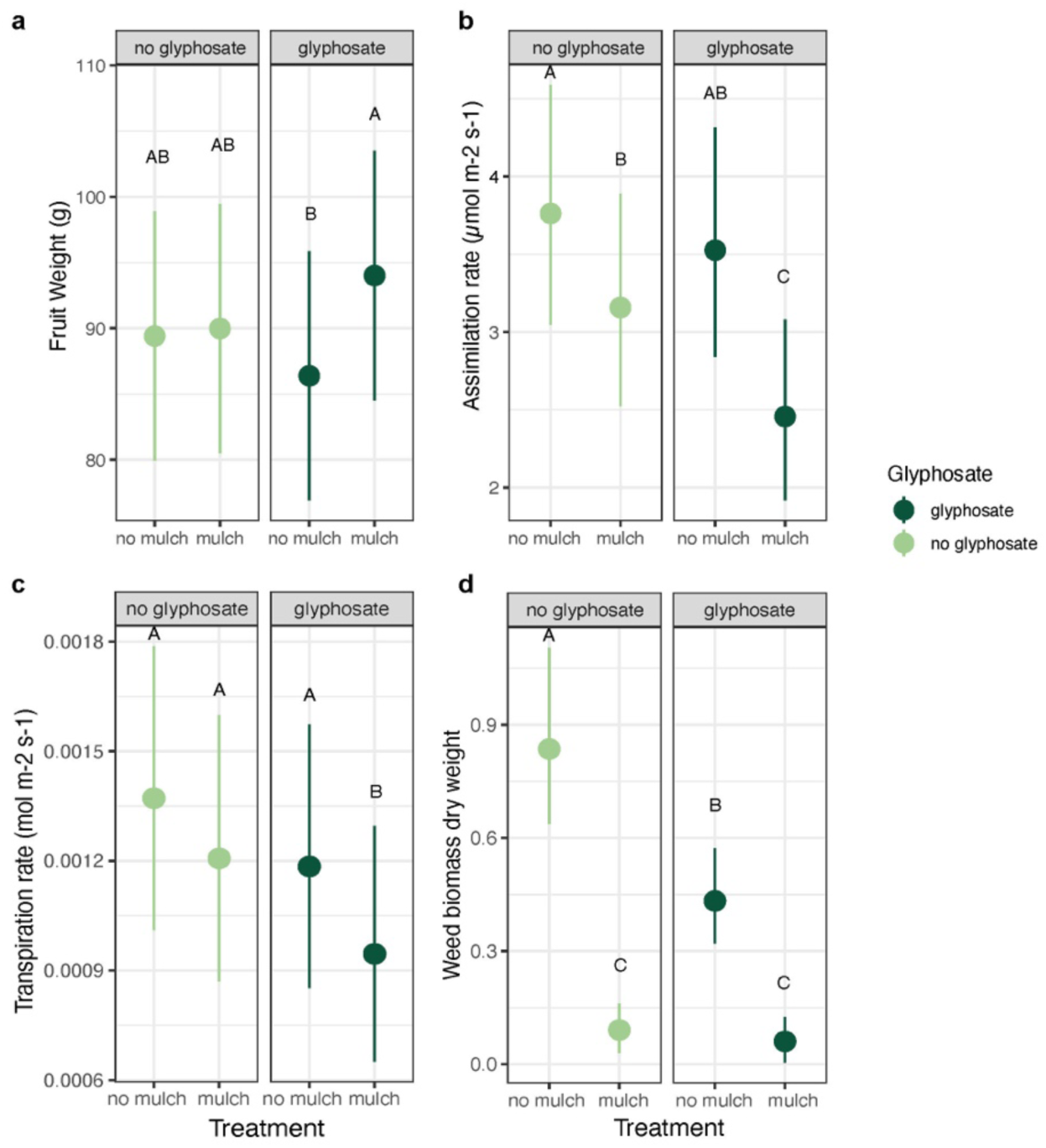
Mulch and glyphosate field applications significantly altered mature tree performance, productivity, and weed biomass. For all plots, points are estimated marginal means of biological replicates, error bars represent standard error, and letters indicate pairwise group differences based on ANOVA followed by a two-tailed Tukey post hoc test and FDR correction (P < 0.05). a, Average fruit weight per tree was measured in grams and includes n=128 tree samples. b, carbon assimilation rate measured in µmol m^-2^ s^-1^ and includes n=192 tree samples across three years. c, Transpiration rate was measured in mol m^-2^ s^-1^ and includes n=192 tree samples across three years. d, Total weed dry weight was measured in kilograms and includes n=48 plots across two years of sampling.

Weed biomass responded strongly to the field applications (Fig. 3d). Mulch significantly reduced weed biomass relative to unmulched plots (ANOVA, F_1,44_=87.193, P=5.29×10^-12^). Glyphosate applied alone also reduced weed biomass, but produced an intermediate phenotype relative to mulch treatments (ANOVA, F_1,44_=12.353, P=0.0010). The combination of mulch and glyphosate did not differ significantly from mulch alone, indicating that mulch was the dominant driver of weed suppression.

### Mulch enriched saprophytes and likely transmitted a putative fungal pathogen

Next, we completed a differential abundance analysis to identify microbial taxa that may be contributing to the changes in tree trait expression. Glyphosate and HA treatments did not cause significant bacterial or fungal genera enrichments or depletions in the root or rhizosphere. Root bacterial and fungal genera enriched in mulched plots included *Zoogloea*, *Candolleomyces*, and *Coniochaeta*, while an unknown genus in the Methylophilaceae family was significantly depleted (Fig. 4a,c). The rhizosphere bacterial and fungal communities had 59 and 27, respectively, differentially abundant genera between mulch-treated and no mulch plots (Fig. 4b,d, Supplemental Table S2). Mirroring results above, this further confirms that the citrus root microbiome was more resilient to abiotic changes than the rhizosphere microbial community.

**Figure 4.**
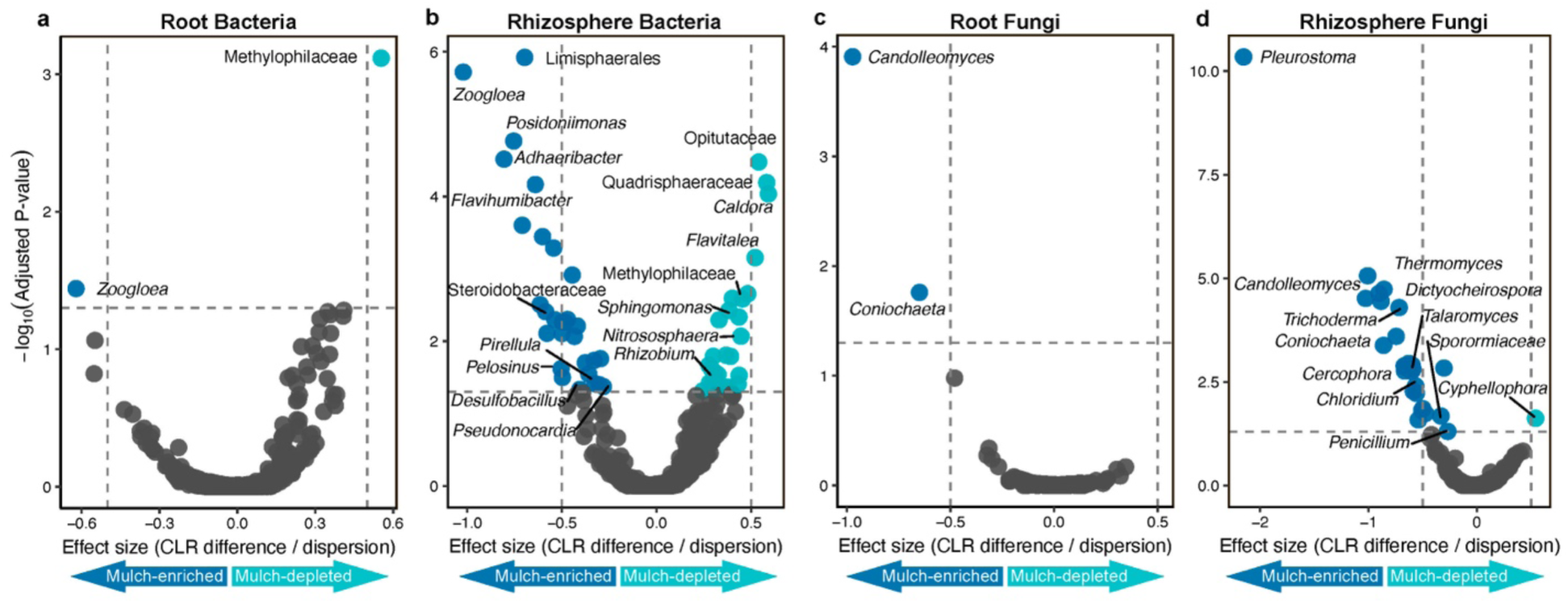
Mulch application significantly enriched for saprophytes and a putative pathogen in the rhizosphere bacterial and fungal communities. The bacterial root (a), bacterial rhizosphere (b), fungal root (c), and fungal rhizosphere (d) plots are volcano plots indicating mulch-enriched (dark blue points) and mulch-depleted (light blue points) genera. Grey points represent genera that are not significantly enriched. Note that a few archaea genera are included in the root and rhizosphere bacterial analyses and plots. If the genus (italicized) name was unknown the order or family name is listed. Enrichments were calculated using n=95 rhizosphere and n=89 root bacterial, and n=95 rhizosphere and n=92 root fungal biological replicate samples. Full statistical analyses are available in Supplemental Table S2.

Among the most strongly enriched taxa in mulch-treated plots were saprotrophic fungi (*Thermomyces, Dictyocheirospora, Trichoderma, Talaromyces, Cercophora, Chloridium,* etc.) [69–74] and bacteria associated with aquatic environments and complex carbon degradation (*Limisphaerales*, *Zoogloea*, *Flavihumibacter*, *Pirellula*, *Desulfobacillus*, *Pelosinus*, etc.) [75–80] (Fig 4, Supplemental Table S2). This supports the alpha diversity results that bacterial community changes were driven more by rare taxa. Moreover, no-mulch plots were enriched in typical rhizosphere-associated bacteria (*Sphingomonas* and *Rhizobium*) [81, 82] and an ammonia-oxidizing archaeal genus, (*Nitrososphaera*) [83] and the root-associated fungus *Cyphellophora [84]* (Fig 4, Supplemental Table S2). This suggests that mulch application disrupted the ambient community. Of particular interest is the significant enrichment of *Pleurostoma* spp. in the fungal rhizosphere community in mulch-treated plots (Fig. 4d, Supplemental Table S2). *Pleurostoma* has five known species that are wood-inhabiting and cause disease in other woody crops, like olive, grapevine, and almond [85–87].

### Management-driven soil microbiota changes altered plant performance in a greenhouse experiment

To test whether management-induced microbiota shifts likely contributed to the field-observed tree phenotypes, we conducted a greenhouse experiment using soil microbiota collected from each treatment plot approximately one year after the end of the field experiment. Soil microbiota were used to inoculate sterile substrate planted with Carrizo citrus rootstock seeds, the same rootstock variety used in the field experiment. For each soil treatment, an autoclaved (heat-killed) control was included to distinguish active microbial from abiotic soil effects.

Seedling establishment differed strongly between active and heat-killed microbiota treatments (Supplemental Fig. S4a). All autoclaved treatments exhibited 100% establishment, whereas active microbiota from mulch-treated and glyphosate x HA plots significantly reduced seedling establishment to 6.3-18.8%.

After 14 weeks, root and shoot traits were measured. Across treatments, seedlings inoculated with autoclaved microbiota developed significantly larger root systems than those receiving active microbiota (Fig. 5a-c; ANOVA, F_1,171_=82.47, P=2.56 x 10^-16^). Within autoclaved treatments, mulch and glyphosate had no effect. In contrast, for plants receiving active microbiota, both mulch (ANOVA, F_1,171_=7.67, P=0.006) and the mulch x glyphosate interaction (ANOVA, F_1,171_=4.029, P=0.046) significantly reduced root biomass (Fig. 5a-c). Seedlings inoculated with active microbiota originating from plots without mulch and without glyphosate produced the largest roots. This indicates that management practices altered active soil microbiota functions in a way that negatively impacted citrus root development.

**Figure 5.**
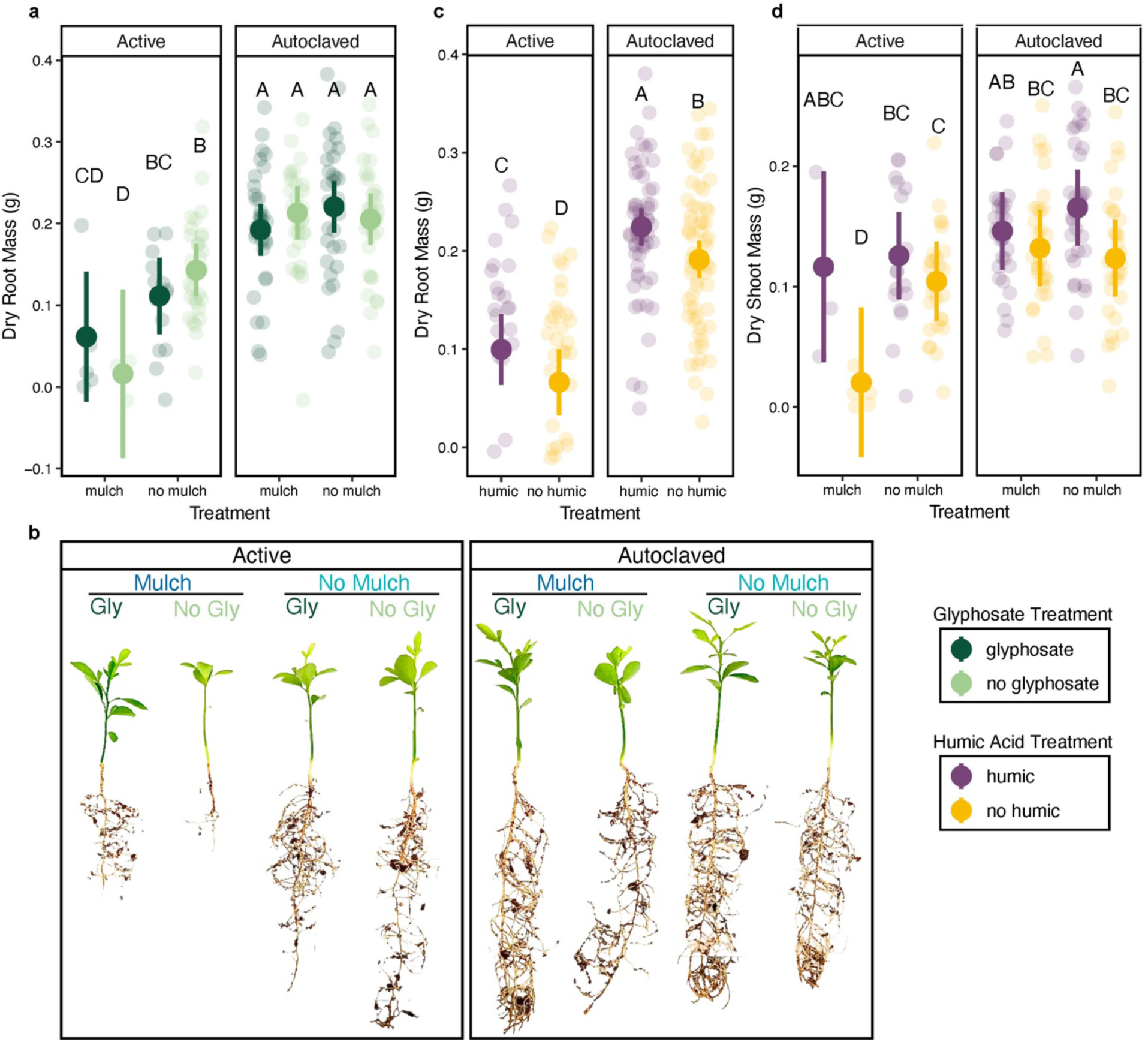
Field applications altered soil microbiota functions that significantly impacted seedling root and shoot mass. For plots a, c, and d, the large opaque points are estimated marginal means of biological replicates, error bars represent upper and lower 95% confidence intervals, and letters indicate pairwise group differences based on ANOVA followed by a two-tailed Tukey post hoc test and FDR correction (P < 0.05). The smaller semi-transparent points represent the raw data measurements of individual biological replicates. These analyses include n=182 biological replicate plant samples. B. includes images of 14 weeks old carrizo rootstock seedlings that are representative of their corresponding treatment group.

Soil originating from HA–treated plots increased root dry mass relative to no HA soils, regardless of whether microbiota were active or heat-killed (Fig. 5c). This suggests that HA primarily influences plant growth directly, rather than through microbial community activity. This compliments our finding that HA had little to no impact on microbiome composition.

Above ground growth responses were limited. Microbiota from glyphosate plots had no effect and were removed from final models. Autoclaved inoculum from HA-treated plots increased aboveground growth. Specifically, seedlings grown with autoclaved soil from HA-treated plots without mulch had significantly greater shoot biomass and height than plants that received autoclaved soil without HA (Fig. 5d; Supplemental Fig. S4b). Active microbiota originating from mulch-treated plots without HA produced significantly smaller shoot weights and heights (Fig. 5d; Supplemental Fig. S4b).

Together, these results indicate that all tested field management practices altered soil microbiota community-level functions in ways that persist even after application ceased and can strongly influence citrus seedling establishment and growth, with potential implications for mature field tree growth and health. The negative effect of active microbiota from mulch treated plots paralleled the negative physiological responses of field trees in mulched plots, which also experienced the largest shifts in root and rhizosphere microbiome structure. This strongly supports the conclusion that the plant-associated microbial communities are large contributors to the changes in field tree phenotype, rather than just the abiotic effects of mulch applications. However, HA appeared to exert additional plant growth effects independent of microbial activity.

## Discussion

Agricultural inputs act as ecological drivers that restructure soil microbial communities and alter plant–microbe interactions underlying crop productivity. Yet most management strategies were developed without considering their influence on soil and plant microbiota structure and function. Here, we show that common orchard practices reshape citrus root and rhizosphere microbiomes in ways that extend beyond compositional change to influence microbial community-level function and plant performance. Wood mulch application emerged as the dominant ecological driver of microbiome assembly, particularly in the rhizosphere. Mulch altered bacterial and fungal community composition, enriched saprophytic and potentially pathogenic taxa, and interacted with glyphosate to produce the strongest negative effects on tree performance. A greenhouse experiment further demonstrated that these management-induced microbial shifts generated microbiome legacy effects that influenced citrus seedling establishment and root development. These findings support our hypothesis that orchard management practices can restructure microbial communities with consequences for crop performance.

Rhizosphere and root microbiomes had differential responses to field treatments. Root microbial alpha diversity remained stable, whereas rhizosphere diversity increased under mulch, with greater bacterial Shannon and fungal invSimpson indices, indicating that management primarily affected rare bacterial and dominant fungal taxa. A Florida citrus study observed similar patterns of increased bacterial rhizosphere alpha diversity due to wood mulch treatment [30]. These responses suggest that host selective forces are stronger in the root compartment and make the root microbiome more resilient [88, 89].

Mulch was consistently the strongest predictor of microbial community composition. Mulched plots were enriched in saprotrophic fungi and bacteria associated with complex carbon degradation, including taxa commonly found in rotting wood [71, 73, 74, 76, 80, 90] and known for lignocellulose degradation [91]. The enrichment of these organisms suggests that wood mulch introduced large quantities of complex organic carbon and altered the rhizosphere nutrients available and resource competition among microbiome members [92, 93].

In contrast, non-mulched soils were enriched with microbial taxa typically associated with plant roots and rhizosphere nutrient cycling, *Sphingomonas*, *Rhizobium*, and *Nitrososphaera*. *Sphingomonas* spp. are commonly associated with plant growth promotion and phytohormone production [81, 94] and *Rhizobium* and *Nitrososphaera* spp. play important roles in nitrogen fixation [83, 95]. However, non-mulched plots also had significantly more weeds because mulch successfully controlled weeds–even outperforming a glyphosate-based herbicide. These herbaceous plants may have enriched these taxa, which potentially supported weed growth, rather than supporting tree crop growth.

Wood mulch may function as a microbial inoculum, transplanting wood-associated taxa into the rhizosphere [96]. The enrichment of taxa typically associated with wood decomposition supports this possibility. Mulch-treated rhizospheres were enriched with *Pleurostoma*, a fungal genus containing wood-inhabiting pathogens of crops, like olive, grapevine, and almond [85–87]. Considering that the initial mulch used in this study was chipped almond wood, it could have harbored and transmitted *Pleurostoma* spp. Although *Pleurostoma* has not previously been known to cause disease in citrus, the mulch-disturbed microbiome may have facilitated opportunistic infection [97]. Reduced photosynthesis rates in field trees and reduced root growth and establishment in the greenhouse experiment supports this, though further study to implicate *Pleurostoma* as a disease-causing agent in citrus is warranted.

A limitation of this study is that irrigation was kept consistent across all field plots, similar to other studies [98]. However, in practice, mulch application is often accompanied with reduced irrigation because wood mulch increases soil water holding capacity. This may have created an environment conducive for pathogens or other performance-reducing microbes. Additionally, the high carbon-to-nitrogen ratio of hardwood mulch may have caused nitrogen immobilization [99]. Adjusting the irrigation time and frequency in mulched blocks or using alternative mulch materials may have resulted in different outcomes and should be evaluated in follow-up studies.

Interactions among management inputs further influenced microbial community structure. Bacterial community composition was significantly shaped by mulch x glyphosate and mulch x HA interactions, whereas fungal communities responded primarily to main effects. This suggests that bacterial communities may be particularly sensitive to interactions between carbon inputs and chemical disturbances, which can alter microbial community assembly in ways that differ from individual treatments [20, 34]. When combined with large carbon inputs from mulch, glyphosate may create novel selection pressures that restructure bacterial communities by altering both resource availability and imposing metabolic constraints [18, 23, 25, 34]. Notably, the interaction between mulch and glyphosate also produced the strongest negative effects on tree performance in the field, suggesting that microbial responses to combined soil amendments may have important implications for plant–microbe interactions.

Consistent with these interaction effects, rhizosphere microbial diversity was highest under combined treatments, likely reflecting increased resource heterogeneity. However, increased diversity did not correspond to improved plant performance [100]. Trees receiving mulch and glyphosate treatments exhibited lower transpiration and carbon assimilation, despite producing significantly higher average fruit weight. While increases in fruit weight can be beneficial from a crop productivity standpoint, this pattern likely reflects a stress response in which trees allocate more resources toward reproduction under adverse conditions [101].

Unlike mulch and glyphosate, HA had relatively limited effects on microbial community composition, but increased root biomass regardless of whether soil microbiota were active or heat-killed in the greenhouse. This indicates that HA primarily affected plant growth through direct mechanisms rather than through microbial community restructuring. Humic substances influence plant growth by modifying nutrient availability, stimulating root development, and altering plant hormone signaling pathways [32]. HAs have been reported to enrich pathogen suppressive microbial taxa [33], but in our system the dominant effects of HA were not microbiome-mediated.

In addition to providing functional evidence that management-induced changes in the microbiome impact plant performance, the greenhouse experiment also shows that historical field management produces microbiome legacy effects [102]. In other words, microbial community composition and function were influenced by field treatments in ways that persist even after applications cease and soil is transferred into a new environment. Such legacy effects are increasingly recognized in soil ecology and can influence plant-microbe interactions, nutrient cycling, and responses to stress [4, 103].

Overall, this study highlights the importance of considering microbial ecology when evaluating agricultural management practices. Our findings demonstrate that orchard management inputs can restructure rhizosphere and root microbial communities and alter plant-microbe interactions. By understanding how management practices shape microbial community composition and function, it may be possible to design cropping management practices that promote beneficial plant-microbe interactions and minimize unintended microbial community disturbances.

## Acknowledgements

The authors would like to thank the staff of the Lindcove Research and Extension Center for providing support throughout the multi-year field experiment.

## Author contributions

MCR, PR, JS, and NG conceived and designed the study. MCR, PR, JS, AE and SLS provided project administration and supervision. NG, TP, RJ, MIH, and PR collected the data and performed the experiments. EM and SLS processed rhizosphere samples. JWW, VETMA, and MIH processed root samples and built sequencing libraries. JWW, NG, TP, RJ, and JS managed the data. NG and RJ processed and analyzed the data. NG and RJ interpreted the results. NG wrote the original draft of the manuscript. All authors edited and approved the final version of the manuscript.

## Conflicts of interest

The authors declare that they have no known competing financial interests or personal relationships that could have appeared to influence the work reported in this paper.

## Funding

This work was supported by the USDA National Institute of Food and Agriculture (NIFA) Emergency Citrus Disease Research and Extension Program (Grant #2020-70029-33202; PI: MCR, co-PIs: PR, JS, SLS, and AE).

## Data availability

The raw amplicon sequencing data supporting these findings have been deposited in the NCBI SRA repository (*To be released upon publication*) and are associated with BioProject PRJNA1450731. Additionally, all field and greenhouse data and code used in the analyses are publicly available on GitHub and archived on Zenodo [https://doi.org/10.5281/zenodo.19390614 (microbiome reads processing); https://doi.org/10.5281/zenodo.19243975 (microbiome, field, and greenhouse analyses)].

## Supplemental Figure and Table Legends

**Supplemental Figure S1. Photographs of field plots with and without wood mulch application.**

**Supplemental Figure S2. Field applications do not significantly impact root microbiome Alpha diversity.** Bacterial Shannon Index (a), fungal Shannon index (b), bacterial inverse Simpson index (c), and fungal inverse Simpson index (d) plots, points are estimated marginal means of biological replicates, error bars represent standard error, and letters indicate pairwise group differences based on ANOVA followed by a two-tailed Tukey post hoc test and FDR correction (P < 0.05). These analyses include n = 89 bacterial and n = 92 fungal rhizosphere microbiome samples.

**Supplemental Figure S3. Chlorosis was observed in mulch-treated plots.**

**Supplemental Figure S4. Field applications alter soil microbiota functions that significantly impact seedling root and shoot mass.** For all plots, the large opaque points are estimated marginal means of biological replicates, error bars represent upper and lower 95% confidence intervals, and letters indicate pairwise group differences based on ANOVA followed by a two-tailed Tukey post hoc test and FDR correction (P < 0.05). The smaller semi-transparent points represent the raw data measurements of individual biological replicates. These analyses include n=182 biological replicate plant samples.

**Supplemental Table S1. Models and statistical results**. The spreadsheet has four tabs, including models and full statistical test results for alpha diversity, beta diversity, field plant physiology measurements, and greenhouse plant analyses.

**Supplemental Table S2. Mulch differentially abundant microbial taxa analysis results.** The spreadsheet includes four tabs showing the ALDEx2 results for rhizosphere fungi, root fungi, rhizosphere bacteria, and root bacteria. Note, that the bacterial analyses also included eight archeal genera that were amplified by the 16S rRNA primers. Analyses were completed at the genus-level ranking. Columns include taxonomic information (Phylum, Class, Order, Family, and Genus), mean centered-log ratio (CLR) abundance across all samples (rab.all), CLR abundance of mulch (rab.win.mulch) and no mulch (rab.win.no) groups, between group differences (diff.btw), within group variation (dif.win), effect size (effect), proportion of Monte Carlo instances overlap (overlap), Welch’s t-test expected (we.ep) and FDR corrected (we.eBH) P-values, Wilcoxon rank test expected (wi.ep) and FDR corrected (wi.eBH) P-values, the log scale of we.eBH (neglog10_q), significance (sig), and the direction of taxa enrichment (direction).

**Supplemental Table S3. Field tree and weed data.**

**Supplemental Table S4. Greenhouse experiment data.**

## Notes

### Competing Interest Statement

The authors have declared no competing interest.

