## Supplemental Information, Figures, and Text for "Orchard management alters citrus root and rhizosphere microbiomes with functional consequences for plant performance"

**This PDF file includes:**

Supplementary Figures S1 to S4

Tables S1 to S4

Supplemental Methods S1 to S3

Supplemental References

#### Supplemental Figures

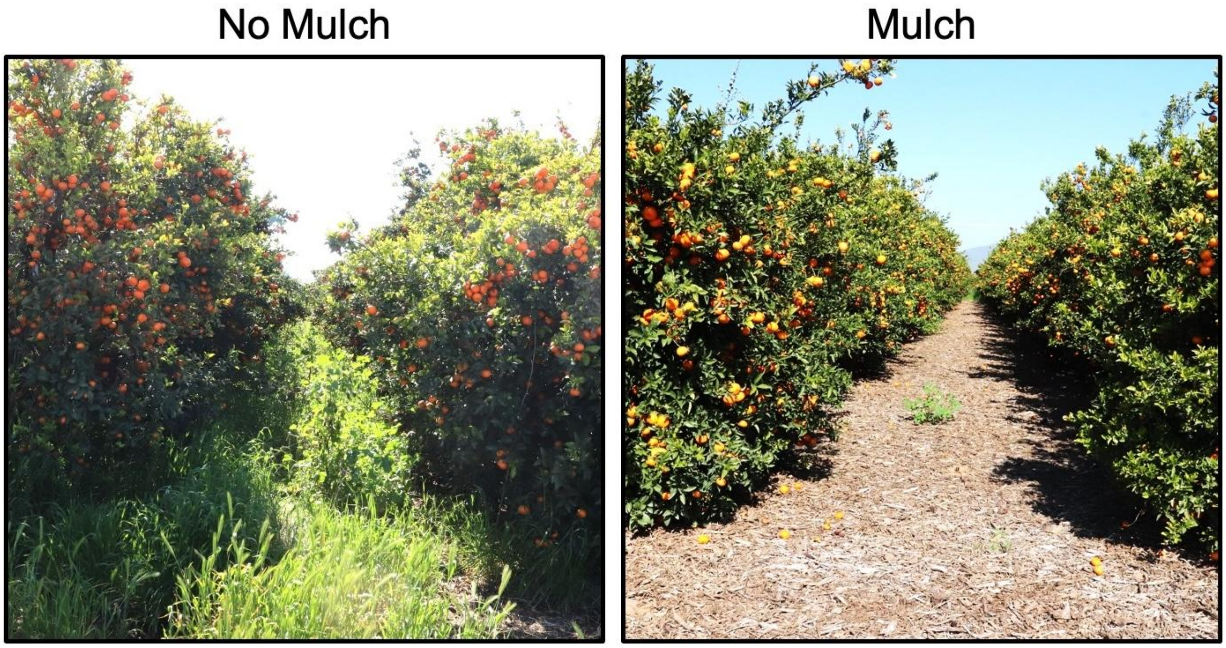


**Supplemental Figure S1. Photographs of field plots with and without wood mulch application.**

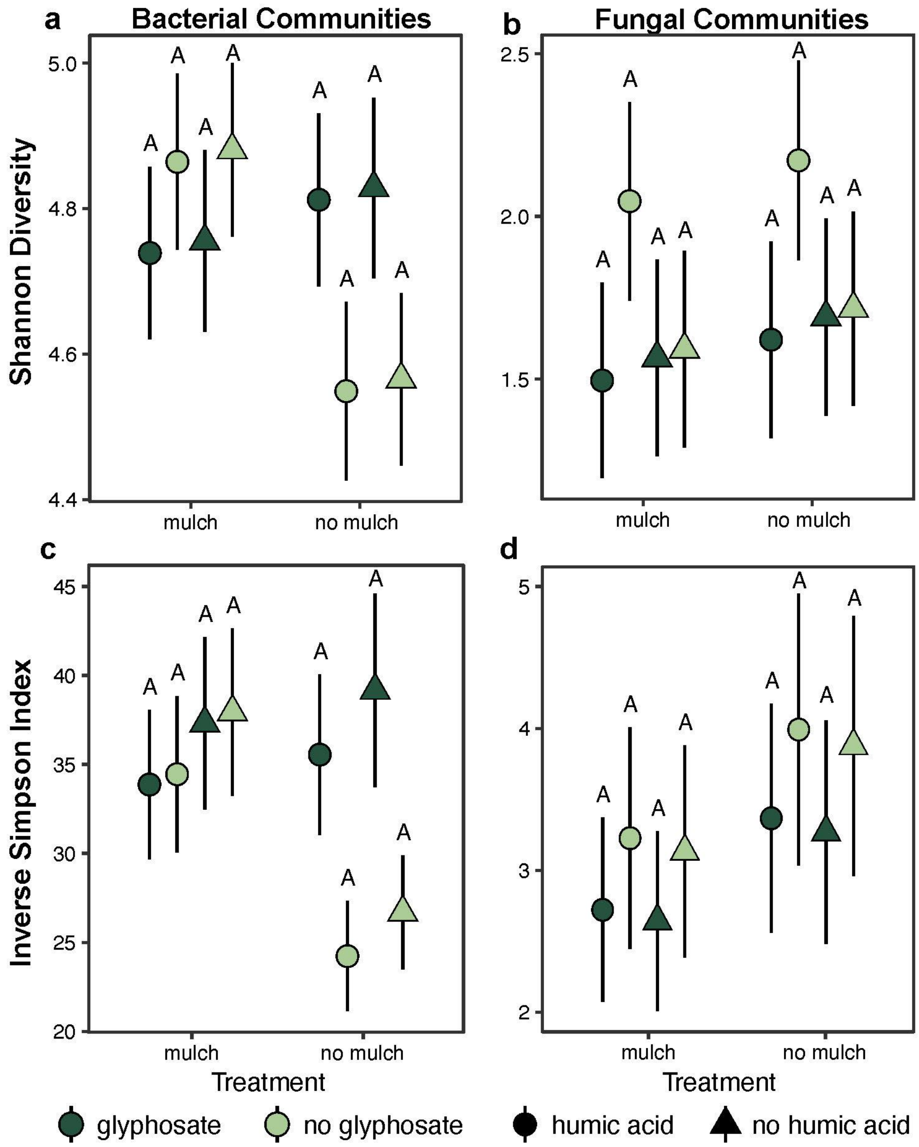


**Supplemental Figure S2. Field applications do not significantly impact root microbiome Alpha diversity.** Bacterial Shannon Index (a), fungal Shannon index (b), bacterial inverse Simpson index (c), and fungal inverse Simpson index (d) plots, points are estimated marginal means of biological replicates, error bars represent standard error, and letters indicate pairwise group differences based on ANOVA followed by a two-tailed Tukey post hoc test and FDR correction ($P < 0.05$). These analyses include n = 89 bacterial and n = 92 fungal rhizosphere microbiome samples.

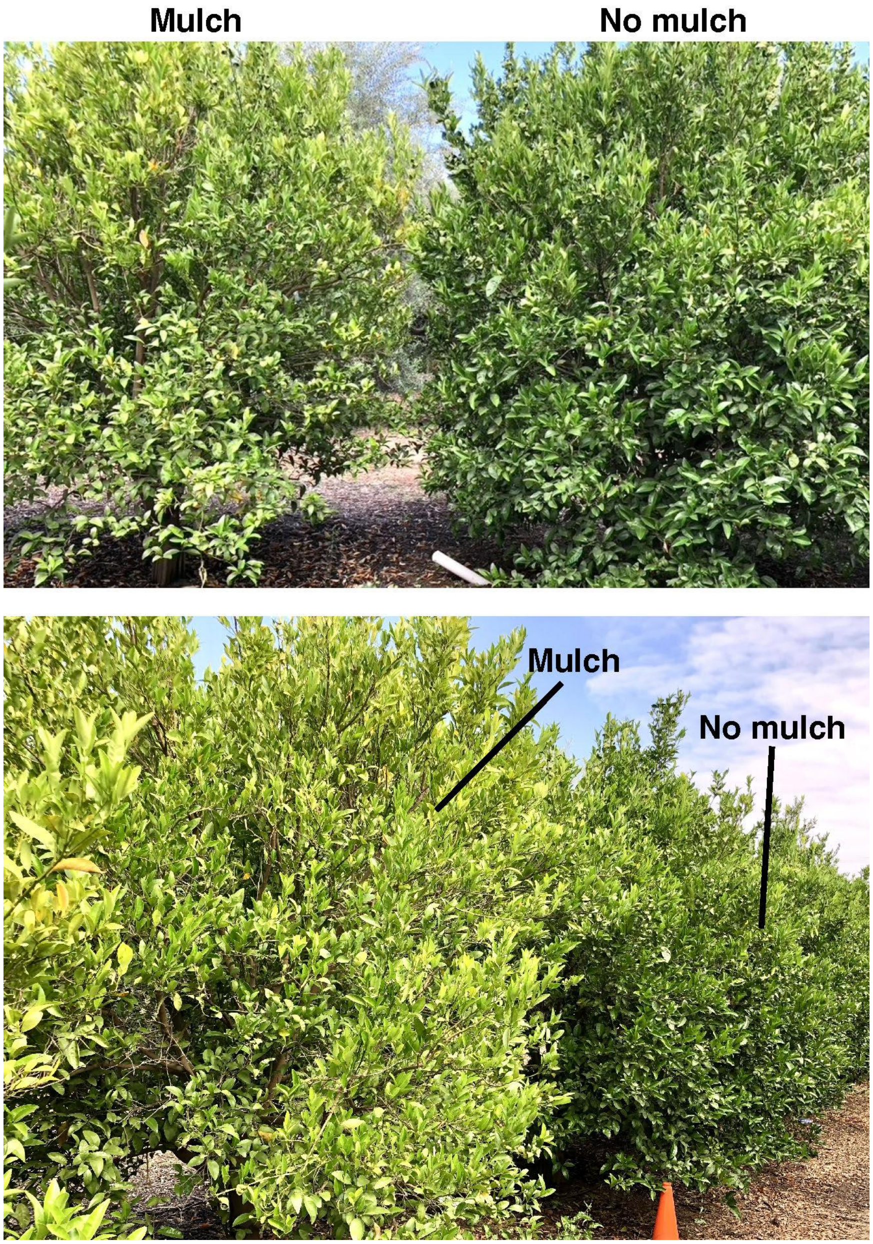


**Supplemental Figure S3. Chlorosis was observed in mulch-treated plots.**

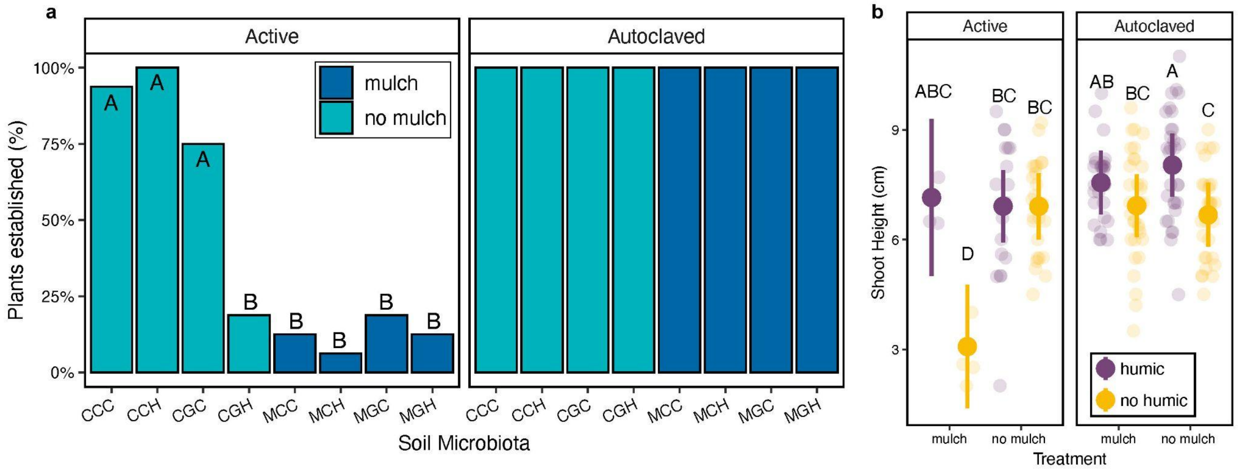


**Supplemental Figure S4. Field applications alter soil microbiota functions that significantly impact seedling root and shoot mass.** For all plots, the large opaque points are estimated marginal means of biological replicates, error bars represent upper and lower 95% confidence intervals, and letters indicate pairwise group differences based on ANOVA followed by a two-tailed Tukey post hoc test and FDR correction ($P < 0.05$). The smaller semi-transparent points represent the raw data measurements of individual biological replicates. These analyses include n=182 biological replicate plant samples.

### Supplemental Tables

**Supplemental Table S1. Models and statistical results**. The spreadsheet has four tabs, including models and full statistical test results for alpha diversity, beta diversity, field plant physiology measurements, and greenhouse plant analyses.
*See supplemental table excel*

**Supplemental Table S2. Mulch differentially abundant microbial taxa analysis results.** The spreadsheet includes four tabs showing the ALDEx2 results for rhizosphere fungi, root fungi, rhizosphere bacteria, and root bacteria. Note, that the bacterial analyses also included eight archeal genera that were amplified by the 16S rRNA primers. Analyses were completed at the genus-level ranking. Columns include taxonomic information (Phylum, Class, Order, Family, and Genus), mean centered-log ratio (CLR) abundance across all samples (rab.all), CLR abundance of mulch (rab.win.mulch) and no mulch (rab.win.no) groups, between group differences (diff.btw), within group variation (dif.win), effect size (effect), proportion of Monte Carlo instances overlap (overlap), Welch’s t-test expected (we.ep) and FDR corrects (we.eBH) P-values, Wilcoxon rank test expected (wi.ep) and FDR corrects (wi.eBH) P-values, the log scale of we.eBH (neglog10_q), significance (sig), and the direction of taxa enrichment (direction).
*See supplemental table excel*

**Supplemental Table S3. Field tree and weed data.**
*See supplemental table excel*

**Supplemental Table S4. Greenhouse experiment data.**
*See supplemental table excel*

### Supplemental Methods

##### *S1. Microbial DNA extractions from field samples.*

Microbial DNA was extracted and purified using a high-throughput workflow implemented on a QIAcube HT robotic instrument (QIAGEN, Hilden, Germany). For citrus root material, 25 mg of lyophilized tissue was added to collection microtubes (#19560, QIAGEN) each containing one 3 mm-diameter tungsten carbide bead (#69997, QIAGEN). The frozen tissue was homogenized using a QIAGEN TissueLyser II for 2 min at 24 Hz in two rounds, first a dry round and then a second round in the presence of 500 µL Solution CD1 (DNeasy 96 PowerSoil Pro QIAcube HT kit, #47021, QIAGEN). The homogenate was centrifuged at 3214 *g* for 15 minutes and the supernatant was combined with 200 µL of Solution CD2 and centrifuged again. The supernatant was transferred to an S-block for DNA purification on the QIACube HT instrument. The *DNeasy 96 PowerSoil Pro for PLANT_QCHT* protocol was modified by replacing Solution CD3 with the viral lysis buffer (#19089, QIAGEN) DNA was eluted in 100 mL of Buffer EB (#19086, QIAGEN).

For rhizosphere sample DNA extraction, samples were defrosted, centrifuged for 8 minutes at 4000g, and 250 mg of the pellet was processed using the DNeasy 96 PowerSoil Pro QIAcube HT Kit (#12855, QIAGEN) following manufacturer's instructions.

Two positive control [75 µL of ZymoBiomics microbial community standard (#D6300, Zymo Research)] and two negative control (sterile Milli-Q water) samples were included in each 96-well plate. DNA concentrations were quantified using the QuantiFluor™ ONE dsDNA System (Promega, Madison, WI, USA) and measured on a BioTek Synergy HTX Multimode Reader (Agilent Technologies, Santa Clara, CA).

##### *S2. Library preparation and amplicon sequencing of bacterial and fungal communities.*

For bacterial amplicon sequencing libraries, 16S V4 rRNA gene amplification was performed using the 515F/806R primers [515F (Parada [1]), GTGYCAGCMGCCGCGGTAA); 806R (Apprill [2]), GGACTACNVGGGTWTCTAAT]. The 515F primers contained 12 basepair unique Golay barcode sequences and both the forward and reverse primers were purchased from IDT (IDT, Coralville, IA, USA) with Illumina adapter sequences attached, according the Earth Microbiome Project protocol [3, 4]. Additionally, Peptide nucleic acid (PNA) oligomers, including pPNA (GGCTCAACCCTGGACAG; PP01-25, PNABio, Thousand Oaks, CA, USA) and mPNA (GGCAAGTGTTCTTCGGA; MP01-25, PNABio, Thousand Oaks, CA, USA) clamps, were included in the PCR to block off-target amplification of citrus plant chloroplast and mitochondria

16S regions [5]. Each PCR reaction contained 10 uL Invitrogen Platinum Hot Start PCR Master Mix (2x) (#13000014, ThermoFisher, Waltham, MA, USA), 3 uL PCR-grade water, 3.75 uL of 5 uM mPNA, 3.75 uL of 5 uM pPNA, 2.0 uL of 2.5 uM 515F, 0.5 uL of 10 uM 806R, and 2.0 uL of template DNA for a total 25 uL volume reaction. For an amplicon size of ~390 bp, the PCRs had an initial denaturation at 94$^o$C for 3 min; followed by 35 cycles of denaturation at 94$^o$C for 45 s, annealing PNA clamps at 78$^o$C for 10 s, annealing the primers at 50$^o$C for 60 s, and extension at 72$^o$C for 90 s; followed by a final extension at 72$^o$C for 10 min. Each 96-well plate PCR included an additional two negative controls (PCR-grade water).

For fungal amplicon sequencing libraries, ITS1 region amplification was completed using the ITS1f/ITS2 primers [6] (ITS1f, CTTGGTCATTTAGAGGAAGTAA); ITS2, GCTGCGTTCTTCATCGATGC) based on the Earth Microbiome Project protocol [4, 7]. The ITS2 reverse primer contained the 12 bp unique Golay barcode sequences and both the forward and reverse primers were purchased from IDT (IDT, Coralville, IA, USA) with Illumina adapter sequences attached. Each PCR reaction contained 10 uL Platinum Hot Start PCR Master Mix (2x) (#13000014, ThermoFisher, Waltham, MA, USA), 10.5 uL PCR-grade water, 0.5 uL of 10 uM ITS1, 2.0 uL of 2.5 uM ITS2, and 2.0 uL of template DNA for a total 25 uL volume reaction. For an amplicon size of ~250-600 bp, the PCRs had an initial denaturation at 94$^o$C for 1 min; followed by 35 cycles of denaturation at 94$^o$C for 30 s, annealing at 52$^o$C for 30 s, and extension at 68$^o$C for 30 s; followed by a final extension at 68$^o$C for 10 min.

Bacterial and fungal PCRs were performed in duplicate and products were confirmed on a 1% agarose gel and then duplicates were pooled. Twenty-five µL of PCR product per sample were purified using the SequalPrep Normalization Plate Kit (ThermoFisher #A1051001) according to the manufacturer's recommendations. Then, 5 µL of each sample from one 96-well plate was pooled into one tube and underwent quality checks (QC) using a tapestation and qPCR at the UC Riverside Genomics Core Facility (UCR GCF). After QC, pooled samples were combined to have a final ratio of 2:3 16S:ITS and concentrated with Ampure XP Beads (Beckman Coulter #A63880) in a 1.8X bead ratio. The final pool followed the same QC pipeline prior to sequencing.

##### *S3. Amplicon sequence processing.*

Raw Illumina sequences were demultiplexed, with zero index barcode mismatches allowed, and converted to FASTQ format using BCL Convert v3.8.4. Adaptors and primer sequences were removed using trimmomatic [8] with standard settings and cutadapt [9] with a minimum 5-base overlap, maximum 15% error rate when matching primers, and used -–

nextseq-trim=20 to trim polyX tails and low-quality ends. FastQC [10] was used to verify read quality and check for adapter sequences. Additional quality control and reads processing was completed using the DADA2 pipeline [11], see full workflow in Supplemental Methods S3.

For 16S reads, forward reads were discarded if they had more than 3 expected errors, otherwise they were truncated at 230 nucleotides; for reverse reads the parameters were 4 expected errors and 210 nucleotides. Error rates were estimated separately for forward and reverse reads based on a sample of the first 500 reads from each sequence file and then used to denoise and de-replicate reads using the standard DADA2 functions. Chimeric sequences were detected and removed using the 'consensus' procedure in DADA2. Each individual sample was processed in parallel, after which all the resulting ASV tables were merged. Finally, the taxonomy of each ASV was assigned using the DADA2 assignTaxonomy() function, which implements a naive Bayesian classifier, and the GreenGenes2 09.2024 release database [12].

For the ITS reads, forward reads were discarded if they had more than 3 expected errors, otherwise they were truncated at 220 nucleotides; for reverse reads the parameters were 4 expected errors and 220 nucleotides. Then, the ITS reads were processed in the same pipeline outlined for the 16S reads above. Finally, fungal taxonomy was assigned using the assignTaxonomy() function and the UNITE v.10 (02.19.2025 release) fungal database [13].
